# The dual role of a multi-heme cytochrome in methanogenesis: MmcA is important for energy conservation and carbon metabolism in *Methanosarcina acetivorans*

**DOI:** 10.1101/2022.07.20.500911

**Authors:** Blake E. Downing, Dinesh Gupta, Dipti D. Nayak

## Abstract

Methanogenic archaea belonging to the Order *Methanosarcinales* conserve energy using an electron transport chain (ETC). In the genetically tractable strain *Methanosarcina acetivorans*, ferredoxin donates electrons to the ETC via the Rnf (**<u>R</u>**hodobacter **<u>n</u>**itrogen **f**ixation) complex. The Rnf complex in *M. acetivorans*, unlike its counterpart in Bacteria, contains a multiheme *c*-type cytochrome (MHC) subunit called MmcA. Early studies hypothesized MmcA is a critical component of Rnf, however recent work posits that the primary role of MmcA is facilitating extracellular electron transport. To explore the physiological role of MmcA, we characterized *M. acetivorans* mutants lacking either the entire Rnf complex (Δ*rnf*) or just the MmcA subunit (Δ*mmcA*). Our data show that MmcA is essential for growth during acetoclastic methanogenesis but neither Rnf nor MmcA are required for methanogenic growth on methylated compounds. On methylated compounds, the absence of MmcA alone leads to a more severe growth defect compared to a Rnf deletion likely due to different strategies for ferredoxin regeneration that arise in each strain. Transcriptomic data suggest that the Δ*mmcA* mutant might regenerate ferredoxin by upregulating the cytosolic Wood-Ljundahl pathway for acetyl-CoA synthesis, whereas the Δ*rnf* mutant may repurpose the F_420_ dehydrogenase complex (Fpo) to regenerate ferredoxin coupled to proton translocation. Beyond energy conservation, the deletion of Rnf or MmcA leads to some shared and some unique transcriptional changes in methyltransferase genes and regulatory proteins. Overall, our study provides systems-level insights into the non-overlapping roles of the Rnf bioenergetic complex and the associated MHC, MmcA.

**Importance:** Methane is a greenhouse gas that is ten times more potent than carbon dioxide and efforts to curb emissions are crucial to meet climate goals. Methane emissions primarily stem from the metabolic activity of microorganisms called methanogenic archaea (methanogens). The electron transport chain (ETC) in methanogens that belong to the Order *Methanosarcinales* has been the focus of many *in vitro* studies to date, but the endogenous functions of the bioenergetic complexes that comprise the ETC have rarely been investigated. In this study, we use genetic techniques to functionally characterize the Rnf bioenergetic complex and the associated multi-heme *c*-type cytochrome MmcA in the model methanogen, *Methanosarcina acetivorans*. Our results show that MmcA and Rnf have shared and unique roles in the cell, and that, contrary to current knowledge, *M. acetivorans* has the capacity to induce at least two alternative pathways for ferredoxin regeneration in the absence of a functional Rnf complex.

## Introduction

The vast majority of methane released to the atmosphere is generated by a group of microorganisms called methanogens (1). Methanogens belong to the Domain Archaea and produce methane as a by-product of energy conservation (2). Methanogens are polyphyletic, and the most widely distributed mode of methanogenic growth uses molecular hydrogen to reduce carbon dioxide to methane (2, 3). Growth on hydrogen and carbon dioxide (termed hydrogenotrophic methanogenesis) is a highly conserved and well-characterized seven-step pathway (Supplementary Figure 1a) (3). A few methanogens, notably members of the Order *Methanosarcinales,* have an expanded metabolic repertoire that includes growth on small organic acids like acetate (via acetoclastic methanogenesis) or on methylated compounds like methanol or methylamines (via methylotrophic methanogenesis) (Supplementary Figure 1b-1e) (2, 4). The metabolic versatility of the *Methanosarcinales* is linked to the presence of a membrane-bound electron transport chain (ETC) for energy conservation, which is absent in many other methanogen lineages (2, 4).

In the *Methanosarcinales*, the ETC can be broken down into one or more input modules, a membrane-bound electron carrier called methanophenazine (MP), and a single output module (4, 5). The input module(s) serve as an entry point for electrons from a variety of electron donors and cofactors into the ETC whereas the output module transfers electrons to the terminal electron acceptor (4). Depending on the strain and the growth substrate, the input module(s) can vary substantially but the output module remains constant because the terminal electron acceptor is always a heterodisulfide (CoM-S-S-CoB) of two cofactors, coenzyme M (CoM) and coenzyme B (CoB) (Figure 1) (3, 4, 6). The CoM-S-S-CoB is generated by the enzyme methyl coenzyme M reductase (MCR) in the last step of methanogenesis (Supplementary Figure 1) (3, 6). Accordingly, all members of the *Methanosarcinales* encode a membrane-associated heterodisulfide reductase complex (HdrDE) that serves as the output module of the ETC (4, 7, 8). HdrDE transfers electrons from reduced MP to CoM-S-S-CoB, regenerating coenzyme M (CoM-SH) and coenzyme B (CoB-SH) (Figure 1) (7, 9). Concomitantly, oxidation of MP by HdrDE releases two protons to the pseudoperiplasmic space via a redox loop mechanism (9). In contrast to the omnipresent HdrDE, the input module(s) of the ETC are diverse and, while resembling many of their bacterial counterparts, these bioenergetic complexes have many unique features whose functional ramifications remain poorly characterized.

**Figure 1.**
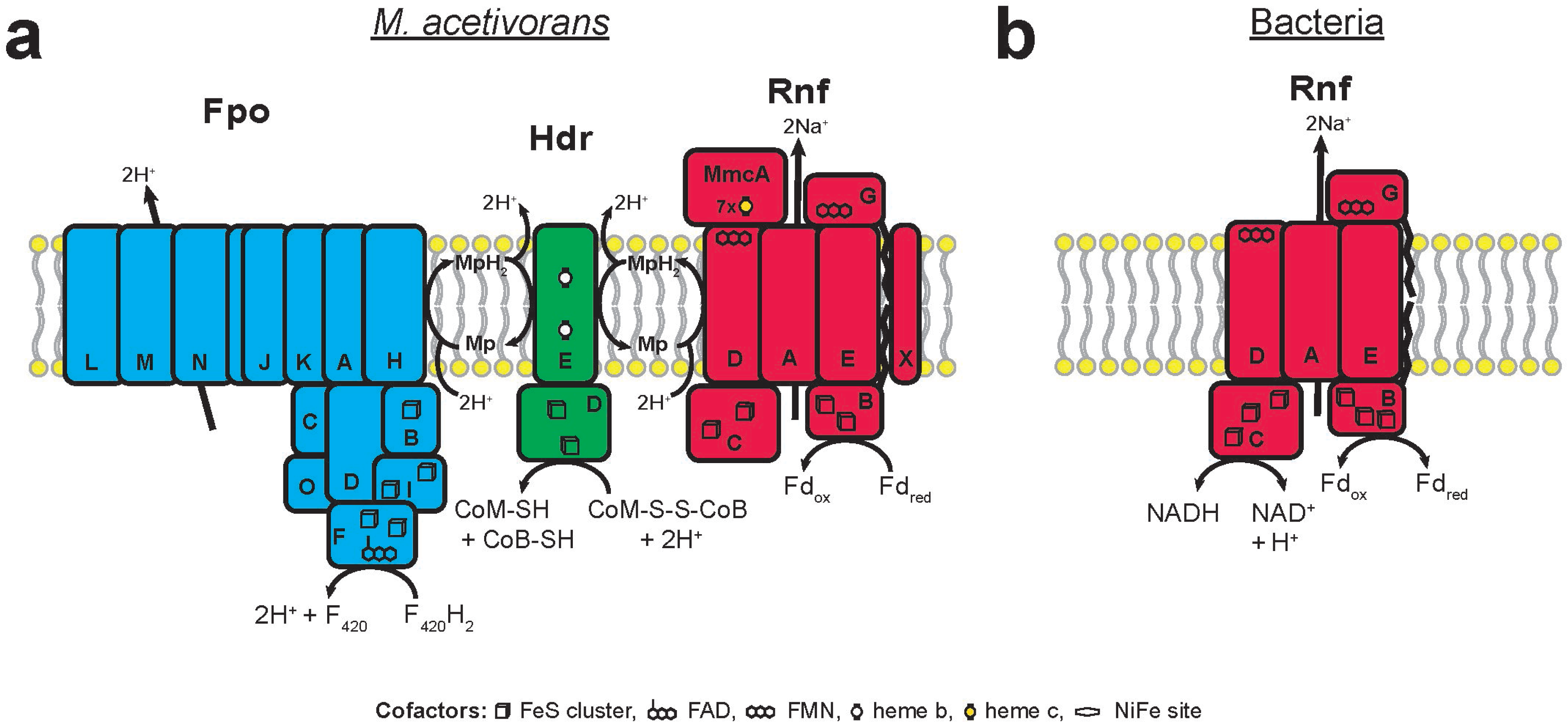
**a)** Schematic of the electron transport chain in *Methanosarcina acetivorans* depicting energy conservation facilitated by two respiratory complexes: 1) the F_420_ dehydrogenase complex in (Fpo) in blue and the *Rhodobacter* nitrogen fixation complex (Rnf) in red along with the terminal electron accepting complex HdrDE in green. **b)** Model of the six-subunit Rnf complex from the bacterium, *Acetobacterium woodii*, which serves as a reversible Na^+^-pumping Ferredoxin (Fd):NAD^+^ oxidoreductase (20). Note: the Rnf complex found in members of the *Methanosarcinales* couples the transfer of electrons between the cytosolic Fd pool and the membrane-bound MP pool to sodium translocation. The exact pathway of electron flow from Fd to MP is unknown. Additionally, in the *Methanosarcinales*, Rnf contains eight subunits instead of six, one of which is a multiheme c-type cytochrome, MmcA.

Within the Genus *Methanosarcina*, strains derived from freshwater environments like *M. barkeri* Fusaro rely on hydrogenases for energy conservation via a process termed “hydrogen cycling” (4, 10–12). However, the marine methanogen *Methanosarcina acetivorans* does not produce any active hydrogenases, and therefore does not rely on hydrogen cycling for the generation of an ion motive force (11, 13). Instead, *M. acetivorans* transfers electrons from reduced cofactors directly to MP via two dedicated bioenergetic complexes: the F_420_ dehydrogenase complex (Fpo) and the *Rhodobacter* nitrogen fixation complex (Rnf) (Figure 1a) (4, 14, 15). The Fpo complex couples the transfer of electrons between the cytosolic F_420_ pool and the membrane-bound MP pool to the translocation of protons across the membrane (Figure 1a) (4, 16). The Fpo complex is related to Respiratory Complex I (RCI) in the mitochondria of eukaryotes and the NADH:ubiquionine oxidoreductase (Nuo) from bacteria, except the NADH interacting module NuoEFG is replaced by the non-orthologous module FpoF (16, 17). In comparison, the Rnf complex couples the transfer of electrons between ferredoxin and MP to the translocation of sodium ions across the membrane (Figure 1a) (15, 18, 19). The genetic organization and cellular function of the Rnf complex in methanogens differs substantially from its bacterial counterparts.

In Bacteria, the Rnf complex is composed of six subunits: RnfABCDEG, and preliminary evidence indicates that electrons flow from reduced Fd to the iron-sulfur clusters of RnfB, then to the covalently bound flavin mononucleotide (FMN) cofactors of RnfG and RnfD, and finally to NAD^+^ in the cytosol via the iron-sulfur clusters of RnfC (Figure 1c) (20). However, no structure has been solved for the complex, so additional cofactors involved in electron flow may be present (20, 21). In addition to the core subunits described above, the *rnf* operon of methanogens also contains an additional gene that encodes a multi-heme *c*-type cytochrome (MHC), *mmcA* (Figure 1d) (18, 22). While the known redox cofactor binding sites of RnfB, RnfD, RnfG and RnfC are conserved in *M. acetivorans,* whether the flow of electrons from Fd from MP follows the same pathway hypothesized in bacterial systems, and how MmcA is involved in this process remains unclear (18, 19). Recent evidence indicates that MmcA in *M. acetivorans* might instead act as a conduit for extracellular electron transfer (EET) to external electron acceptors like anthraquinone-2,6-disulfonate (AQDS), which would substantially broaden the metabolic repertoire of *M. acetivorans* beyond methanogenic growth (22). While it is abundantly clear that MmcA plays an important role in the energy metabolism of *M. acetivorans* and other members of the *Methanosarcinales,* the underlying mechanism(s) remains poorly characterized.

In this study, we use a genetic approach to elucidate the *in vivo* function of MmcA in the model methanogen *M. acetivorans* by comparing the growth and transcriptional responses of mutant strains lacking the MmcA subunit or the entire Rnf complex. Our results show that MmcA might have a cellular function beyond facilitating electron flow through the Rnf complex. Our transcriptomic data also shed light on the coupling between substrate-specific methyltransferases and energy conservation pathways in *M. acetivorans* and bring to light alternate routes for the regeneration of reduced ferredoxin in the absence of Rnf. Overall, our work underscores the importance of MmcA in the methanogen Rnf complex and elucidates a broader role for this multiheme cytochrome during methanogenesis.

## Results and Discussion

### Validation of mutant strains lacking *mmcA* or *rnf* in *Methanosarcina acetivorans*

The *rnf* locus in *M. acetivorans* consists of eight ORFs (MA0658 to MA0665) that encode MmcA and RnfCDGEABX respectively (Figure 2a). The first seven genes have overlapping coding regions and previous studies have also shown that all eight genes are transcribed in a single operon (Figure 2a) (18). To characterize the *in vivo* function of the Rnf complex, we obtained a mutant that has a markerless in-frame deletion in the *M. acetivorans* chromosome spanning *mmcA* through *rnfX* as described in (23) (Figure 2b). To understand the function of the MmcA gene in the Rnf complex, we generated a markerless in-frame deletion in the *mmcA* gene as described previously using Cas9-mediated genome editing (24, 25) (Figure 2c). We sequenced the genome of the Δ*mmcA* mutant and, relative to the parent strain (WWM60), did not detect mutations elsewhere on the chromosome (Supplementary Table 1). Based on this evidence, we can conclude that our MmcA deletion strain does not have any off-target mutations as a result of Cas9 editing or any suppressor mutations to compensate for the loss of *mmcA*.

**Figure 2.**
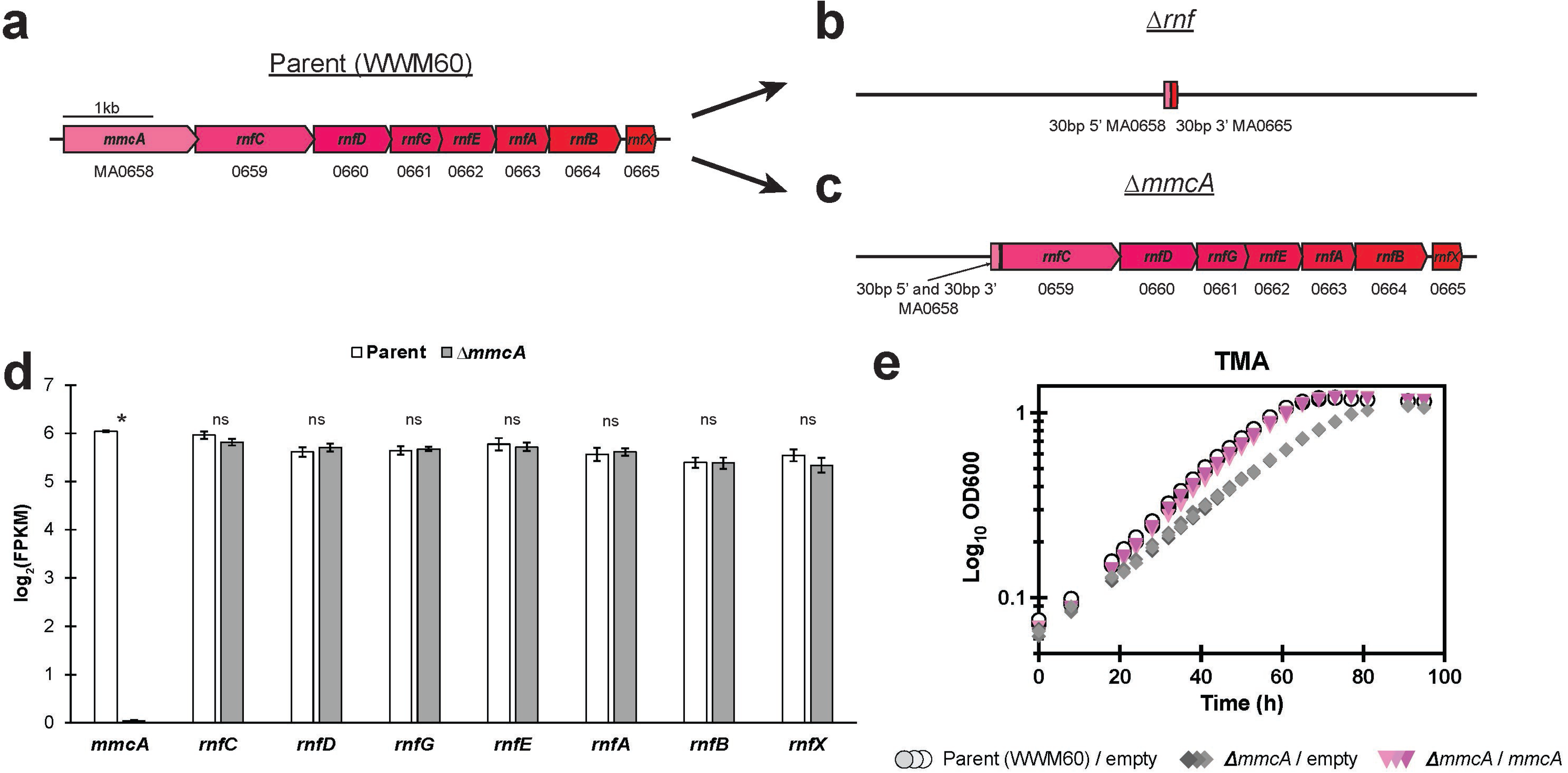
**a)** Chromosomal organization of the eight-gene *rnf* operon in *M. acetivorans*, reflecting the genotype of the parent strain (WWM60) used in this study. Locus tags are provided under each gene. **b)** Chromosomal organization the *rnf* locus in the Δ*rnf* mutant showing a clean deletion of the operon. A 30 basepair (bp) region of the 5’ end of *mmcA* and a 30 bp region of at the 3’ end of *rnfX* are maintained on the chromosome to preclude interference with up- and down-stream regulatory elements outside of the operon. **c)** Chromosomal organization of the *rnf* locus in the Δ*mmcA* mutant. Here, 30 bp regions at the 5’ and 3’ ends of *mmcA* are maintained to prevent frameshift disruption of downstream *rnf* genes. **d)** Expression values for the *rnf* genes in the parent and Δ*mmcA* strains. Expression is measured in log_2_(FPKM) [fragments per kilobase of transcript per million mapped reads] for each gene. In the Δ*mmcA* strain, only *mmcA* expression is abolished [p-value = 3.5E-9; Welch’s t-test] indicating the deletion does not adversely impact on the expression of other genes in the operon. Significance (p-values < 0.05) for log_2_(FPKM) values of individual genes between strains is indicated with an asterisk or “ns” for non-significant change. **e)** Growth curve of the parent strain carrying an empty vector control (light gray circles), the Δ*mmcA* mutant carrying an empty vector control (dark gray diamonds), and the Δ*mmcA* mutant carrying a plasmid expressing *mmcA* (pink inverted triangles) in HS medium containing 50 mM TMA as the sole carbon and energy source. The empty vector contains the β-glucuronidase gene, *uidA*, under the control of a tetracycline inducible P*mcrB* (*tetO4*) promoter described previously in (60). The complementation vector contains *mmcA* under the control of a tetracycline inducible P*mcrB* (*tetO4*) promoter described previously in (24). Three replicate tubes of each strain were used for growth assays and tetracycline was added to a final concentration of 100 ug/mL in the growth medium.

Since *mmcA* is the first gene in the *rnf* operon, it is likely that deletion of *mmcA* could alter the expression of the other *rnf* genes. To test for polar effects, we measured the expression of *rnfCDGEABX* in the Δ*mmcA* mutant and WWM60 using whole-genome RNA sequencing during growth on trimethylamine (TMA) and did not detect significant change in transcript levels of any of these genes in the absence of *mmcA* (p-value >0.05; Welch’s t-test) (Figure 1d). Thus, knocking out the *mmcA* gene does not alter the transcription of *rnfCDGEABX*. In addition, we observed that expression of *mmcA in trans* functionally complements the chromosomal deletion of *mmcA* and restores wildtype growth on TMA (Figure 2e). These growth data indicate that the RnfCDGEABX proteins are produced in the Δ*mmcA* mutant and that expression of *mmcA* in *trans* can reconstitute a functional Rnf complex.

### Growth characteristics of *rnf* and *mmcA* mutants during methanogenesis on different substrates

*In vitro* analyses of the Rnf complex from *M. acetivorans* suggest that MmcA plays an important role in mediating electron flow between ferredoxin and MP, but the importance of MmcA for Rnf activity *in vivo*, outside of extracellular electron transfer, has not been well characterized (22). To address this gap in knowledge, we measured the growth phenotype of WWM60, Δ*rnf*, and Δ*mmcA* on a wide range of substrates that represent the metabolic breath of *M. acetivorans*.

The Rnf complex as well as MmcA are essential for growth on acetate via acetoclastic methanogenesis (18, 23, 24). We observed no growth of the Δ*rnf* or Δ*mmcA* mutants on acetate, while WWM60 with a fully functional Rnf complex was viable (Figure 3a, Table 1). We attempted to isolate suppressor mutants in the Δ*rnf* or Δ*mmcA* backgrounds that restore growth on acetate by incubating each strain in growth medium containing acetate as the sole source of carbon and energy. We did not detect any observable growth (measured as a change in optical density) for all three replicates of either strain after an incubation period of one year. Together, these results indicate that the Rnf complex is essential during methanogenesis from acetate for regenerating reduced ferredoxin generated by the carbon monoxide dehydrogenase/acetyl CoA synthase (CODH/ACS) complex (18, 26, 27). In addition, the MHC subunit MmcA is vital for the functionality of the Rnf complex. Our growth data for the Δ*mmcA* mutant on acetate agree with previous work from our group, but are in contrast a previous study where a *mmcA* deletion mutant of *M. acetivorans* had no growth defect on acetate compared to the corresponding parent strain (22, 24). We suspect that the difference in phenotype observed may pertain to differences in the genetic techniques used for generating deletion mutants, or the presence of a suppressor mutation in the *mmcA* deletion strain generated in the previous work (22).

**Figure 3.**
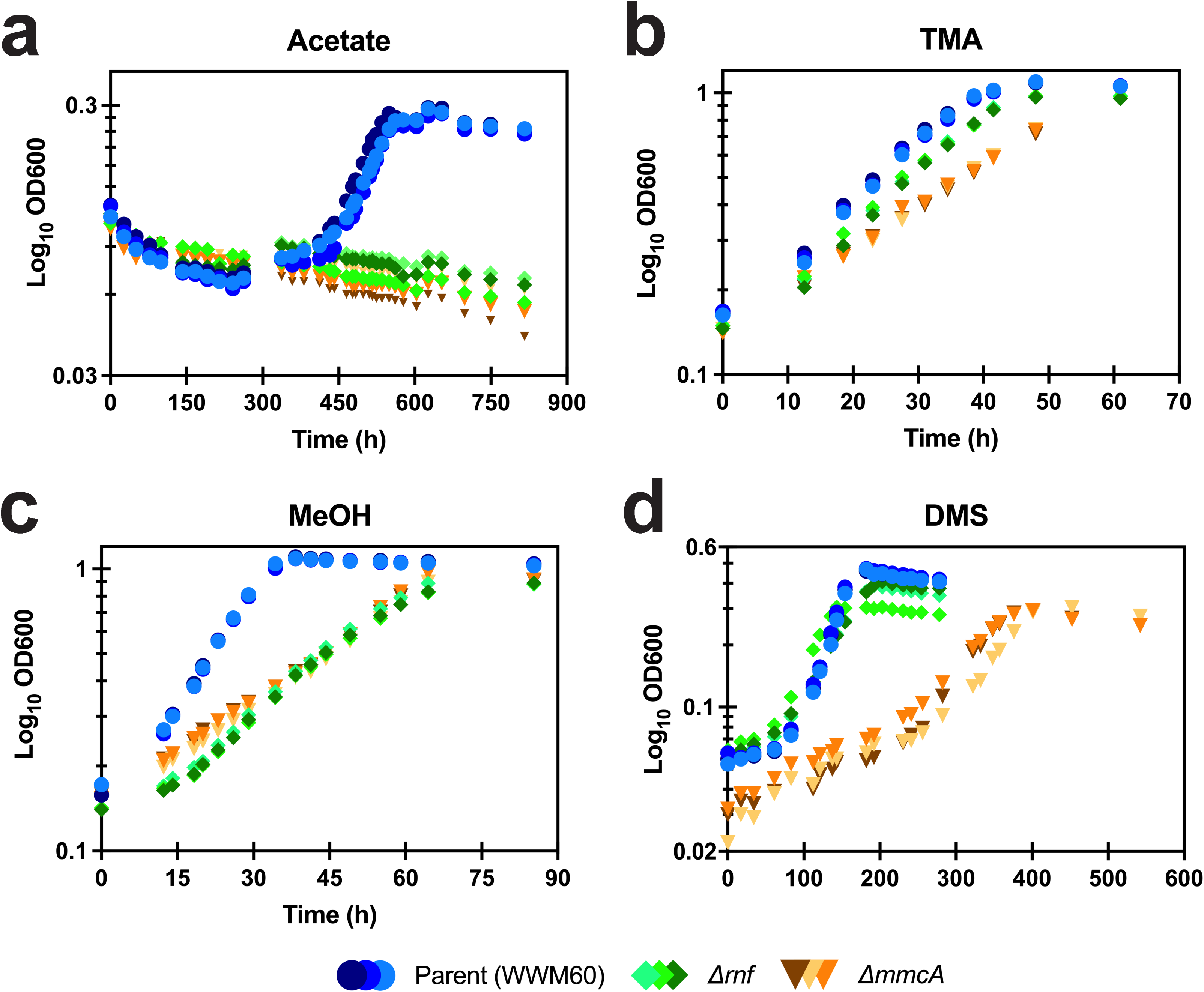
Growth curves of the parent strain (WWM60) (blue circles), *Δrnf* (green diamonds), and Δ*mmcA* (orange inverted triangles) mutants on **a)** 40 mM acetate, **b)** 50 mM trimethylamine (TMA), **c)** 125 mM MeOH, and **d)** 20 mM dimethylsulfide (DMS) as the sole carbon and energy source. Three replicate tubes of each strain were used for growth assays.

**Table 1:** Growth data for WWM60, WWM1015 (Δrnf) and DDN009 (ΔmmcA) on a range of different methanogenic substrates

| *M. acetivorans* strains | Doubling time (Td) (hrs) | Max.OD<sub>600</sub> (OD<sup>M</sup>) | p-value for WT vs $\Delta rnf$ mutant (Td/OD<sup>M</sup>) | p-value for WT vs $\Delta mmcA$ mutant (Td/OD<sup>M</sup>) | p-value for $\Delta rnf$ vs $\Delta mmcA$ mutant (Td/OD<sup>M</sup>) |
| --- | --- | --- | --- | --- | --- |
| 50 mM Trimethylamine (TMA) as a growth substrate |  |  |  |  |  |
| WWM60 (WT) | 14.94 ± 0.61 | 1.698 | 0.043/0.035 | 0.001/0.355 | 0.009/0.063 |
| $\Delta rnf$ | 16.72 ± 0.86 | 1.485 | - | - | - |
| $\Delta mmcA$ | 20.07 ± 0.40 | 1.632 | - | - | - |
| 125 mM Methanol (MeOH) as a growth substrate |  |  |  |  |  |
| WWM60 (WT) | 10.33 ± 0.23 | 1.899 | 8.7616E-06/0.001 | 9.0135E-06/0.002 | 0.001/0.783 |
| $\Delta rnf$ | 18.58 ± 0.11 | 1.558 | - | - | - |
| $\Delta mmcA$ | 25.09 ± 0.36 | 1.569 | - | - | - |
| 40 mM Acetate as a growth substrate |  |  |  |  |  |
| WWM60 (WT) | 79.48 ± 5.73 | - | - | - | - |
| $\Delta rnf$ | No growth | - | - | - | - |
| $\Delta mmcA$ | No growth | - | - | - | - |
| 20 mM Dimethyl sulfide (DMS) as a growth substrate |  |  |  |  |  |
| WWM60 (WT) | 26.48 ± 0.13 | - | 0.023 | 0.016 | 0.057 |
| $\Delta rnf$ | 48.54 ± 5.90 | - | - | - | - |
| $\Delta mmcA$ | 67.14 ± 8.90 | - | - | - | - |

All data represent the mean ± standard deviation of at least 3 biological replicates p-values were determined using a two-sided t-test assuming unequal variances

Next, we tested the role of Rnf and MmcA during methylotrophic methanogenesis by measuring the growth phenotype of the mutants and the parent strain on three different methylated compounds: trimethylamine (TMA), methanol (MeOH), and dimethylsulfide (DMS) (14, 28–30). The Δ*rnf* mutant and the Δ*mmcA* mutant were viable on all three methylated compounds, but we noted a significant defect in the growth rate of both mutants relative to WWM60 (Figure 3b, Table 1). Additionally, the *ΔmmcA* mutant had a significantly higher growth defect compared to the Δ*rnf* mutant on all methylated compounds (Figure 3; Table 1). Based on these data, we can conclude that tMmcA and other components of the Rnf complex are not essential for methylotrophic methanogenesis in *M. acetivorans* but are important for optimal growth under these conditions.

The phenotypes of the Δ*rnf* and Δ*mmcA* strains also differed when switching between methylotrophic substrates. We observed that the Δ*rnf* mutant had a significantly shorter lag time switching from medium containing TMA to medium with DMS relative to both WWM60 and the Δ*mmcA* mutant (Figure 4a; Table 2). The lag time of the Δ*mmcA* mutant is not significantly different from the WT strain (Figure 4a; Table 2), further supporting the hypothesis that Rnf proteins are expressed in the absence of MmcA during methylotrophic growth. Based on these data, we hypothesize that loss of the Rnf complex, but not MmcA alone, might result in a transcriptional response tuning the expression of methyltransferases (such as *mtsD*, *mtsF,* and *mtsH*) that are required for growth on methylated sulfur compounds like DMS (30). However, the sensory and regulatory pathway(s) by which this transcriptional response is carried out cannot be inferred from these data.

**Figure 4.**
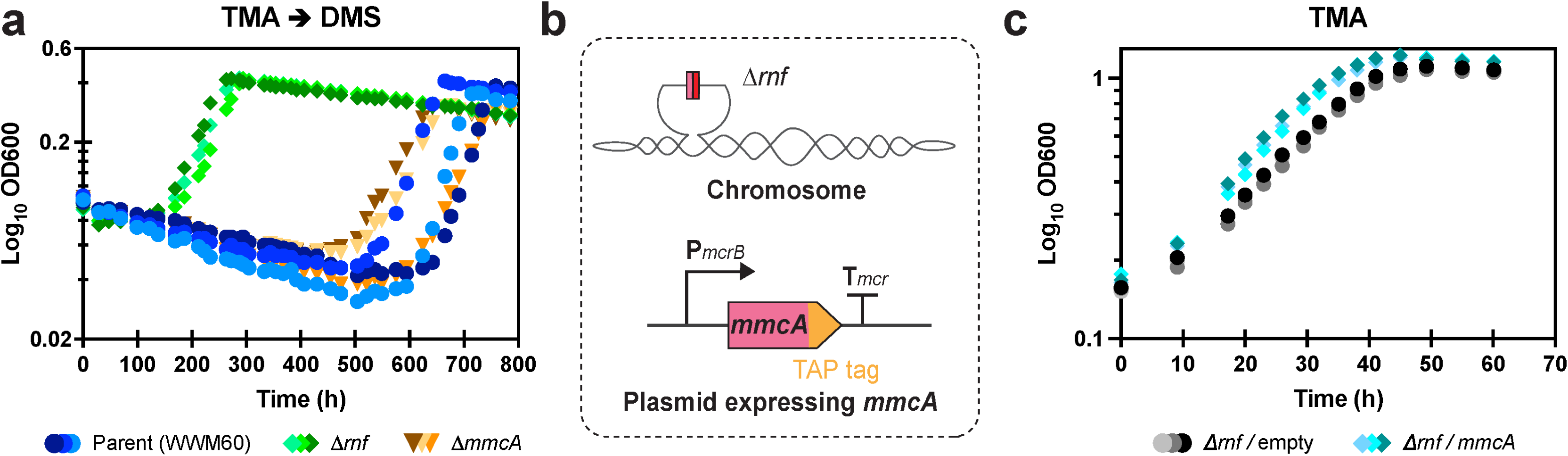
**a)** Growth curve of the parent strain (WWM60) (blue circles), *Δrnf* (green diamonds), and Δ*mmcA* (orange inverted triangles) mutants in HS medium containing 20 mM dimethylsulfide (DMS) as the sole carbon and energy source after transfer from HS medium with 50 mM trimethylamine (TMA) as the sole carbon and energy source. **b)** A schematic depicting the genotype of a strain generated to assay the role of *mmcA* in the absence of the rest of the *rnf* genes. On the chromosome, the entire *rnf* locus has been deleted. The strain carries an autonomously replicating plasmid encoding *mmcA* with a tandem-affinity-purification (TAP) tag at the C-terminus under the control of a P*mcrB* promoter described previously in (60). **c)** Growth curve of the Δ*rnf* strain carrying an empty vector control (dark gray circles) or a plasmid expressing *mmcA* (light blue diamonds) in HS medium containing 50 mM TMA as the sole carbon and energy source. The empty vector contains the β-glucuronidase gene, *uidA*, under the control of a P*mcrB* promoter described previously in (60). Three replicate tubes of each strain were used for growth assays.

**Table 2:** Growth data for WWM60, WWM1015 (Δrnf) and DDN009 (ΔmmcA) on dimethylsulfide (DMS)

| *M. acetivorans*<br>strains | Doubling<br>time (Td)<br>(hrs) | Lag time<br>(hrs) | p-value for WT vs<br>$\Delta rnf$ mutant<br>(lag time) | p-value for WT vs<br>$\Delta mmcA$ mutant<br>(lag time) | p-value for $\Delta rnf$<br>vs $\Delta mmcA$ mutant<br>(lag time) |
| --- | --- | --- | --- | --- | --- |
| Growth data for cells transferred from TMA to DMS |  |  |  |  |  |
| WWM60<br>(WT) | 36.37 ± 4.52 | 640.60 ± 51.61 | 0.004 | 0.367 | 0.009 |
| $\Delta rnf$ | 53.16 ± 1.92 | 163.42 ± 13.91 | - | - | - |
| $\Delta mmcA$ | 58.68 ± 5.91 | 590.22 ± 68.66 | - | - | - |

All data represent the mean ± standard deviation of at least 3 biological replicates

As the growth defect for the Δ*mmcA* mutant on methylated compounds was significantly exacerbated compared to the Δ*rnf* mutant (Figure 3), it is likely that, in addition to being an important part of the Rnf complex in methanogens, MmcA may have other physiological roles too. To test for a Rnf-independent function of MmcA during methylotrophic methanogenesis, we generated a strain encoding the *mmcA* gene on a plasmid in the Δ*rnf* background (Figure 4b) and measured the growth of this mutant on TMA relative to a control strain carrying an empty vector. Indeed, expression of *mmcA* in the Δ*rnf* background enhanced growth rate compared to the control strain by 22% [p-value = 0.0023; Welch’s t-test] (Figure 4b and Supplementary Table 2). Growth yield of the Δ*rnf* strain expressing *mmcA* was not significantly different than the control strain (Supplementary Table 2). These phenotypic data provide clear evidence that the physiological role of MmcA during methanogenesis from methylated compounds extends beyond its capacity to relaying electrons through the Rnf complex.

### Differential expression of Fpo and methylamine methyltransferases lead to differential growth of the Δ*rnf* and Δ*mmcA* mutants

Based on our growth data, it is clear that the cellular function of the Rnf complex does not completely overlap with the MHC subunit, MmcA, during methylotrophic growth. To understand the genetic basis for this phenotypic distinction, we performed whole genome RNA sequencing of the Δ*rnf* mutant, the Δ*mmcA* mutant and WWM60 at mid-exponential growth on TMA. To identify genes that are “differentially expressed”, we used the following the criteria: a log_2_-fold change in transcript abundance ≥2 (4-fold) with a q-value (corrected p-value) ≤0.01 (see Materials and Methods). The Δ*rnf* mutant had significantly higher expression of genes involved in two distinct aspects of methanogenic metabolism relative to the Δ*mmcA* mutant (Figure 5a). First, we observed significantly higher transcripts for several genes encoding the F_420_ methanophenazine oxidoreductase (Fpo) bioenergetic complex. The membrane bound Fpo complex is comprised of thirteen subunits and, during methylotrophic growth, catalyzes the transfer of electrons from reduced F_420_ to MP coupled to the translocation of two protons across the membrane (Figure 1) (16). Most of the genes comprising the Fpo complex are encoded in the *fpoABCDHIJ1J2KLMNO* operon (Figure 5b) (16, 31). The F_420_ input module *fpoF* is found elsewhere on the chromosome in putative operon with the F_420_-dependent N(5),N(10) -methylene tetrahydromethanopterin reductase (*mer*), and a second copy of *fpoO2* is encoded close to the *fpoA-O* operon, although neither paralog has a known function (Figure 5b) (31). Five genes in the *fpo* operon (*fpoJ2, fpoL, fpoM, fpoN,* and *fpoO*) as well as *fpoO2* had ≥4-fold (or 2 log_2_-fold) higher expression in the Δ*rnf* background compared to the Δ*mmcA* mutant (Figure 5b; Supplementary Table 3). No significant change in expression was observed for *fpoF* (Supplementary Table 3). Thus, we hypothesize that a “headless” or modified form of Fpo that lacks the F_420_ interacting module FpoF is more abundant in the Δ*rnf* mutant. The “headless” Fpo complex has been hypothesized to function as a ferredoxin: MP oxidoreductase in other methanogens, like members of the Genus *Methanothrix (*previously known as Genus *Methanosaeta*) that lack both the Ech hydrogenase as well as the Rnf complex (32). While previous studies have suggested that FpoO plays a role in transferring electrons from the Fpo complex to the MP pool via a [2Fe-2S] cluster, we hypothesize that the FpoO subunit might instead interact with the FpoF subunit and/or with ferredoxin directly (31). It is possible that the two copies of FpoO encoded in the *M. acetivorans* genome differ in their affinity for FpoF versus ferredoxin, which might further modulate the specificity of the Fpo complex for different electron carriers. Altogether, we postulate that higher expression of other subunits of the Fpo complex relative to FpoF and tuning the amount of FpoO2 relative to FpoO1 increases the proportion of the “headless” Fpo complex in the Δ*rnf* mutant (Supplementary Figure 2). The “headless” Fpo complex might provide an alternate route for ferredoxin regeneration in the Δ*rnf* mutant that ultimately leads to faster growth under methylotrophic conditions compared to the Δ*mmcA* mutant. The RNA sequencing data do not allow us to discriminate whether the Fpo genes are upregulated in the Δ*rnf* mutant or downregulated in the Δ*mmcA* mutant. To distinguish between these two modes of regulation, we compared the expression of the Fpo locus in each mutant relative to WWM60. The Fpo genes were more highly expressed in the Δ*rnf* mutant, but not in the Δ*mmcA* mutant, compared to WWM60, however the change in expression for both mutants did not meet our ≥2 log_2_-fold threshold value. Thus, our data suggest the loss of Rnf might result in upregulation of Fpo (Supplementary Table 3), but we cannot confidently identify a mode of regulation that causes a differential expression of the Fpo genes in the Δ*rnf* strain relative to the Δ*mmcA* strain at present.

**Figure 5.**
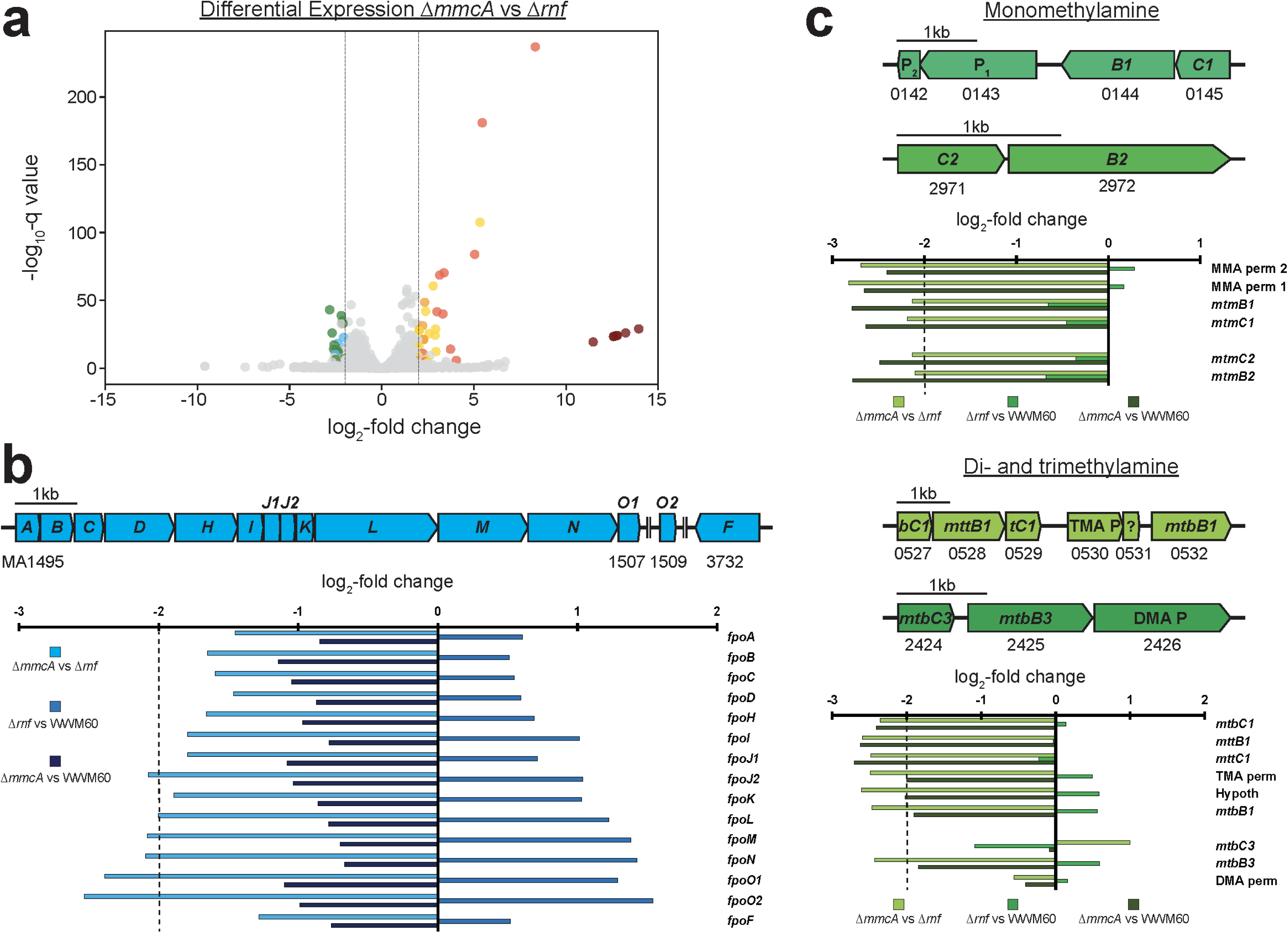
**a)** Volcano plot showing the differential expression of genes between the Δ*mmcA* and Δ*rnf* mutants. Genes with higher expression in the Δ*mmcA* mutant have a positive log_2_-fold change value, while genes with higher expression in the Δ*rnf* mutant have a negative log_2_-fold change value. Dashed lines on the plot delineate the cutoff for ‘significant’ log_2_-fold change in transcript abundance in either mutant. In the Δ*rnf* mutant, two sets of genes are significantly more highly expressed: six subunits of the Fpo complex (shaded in blue), and multiple methylamine methyltransferases and permeases (shaded in green). Genes in orange have higher transcript levels in the Δ*mmcA* mutant relative to both the Δ*rnf* mutant and the parent strain (WWM60), genes in gold have significantly lower transcript levels in the Δ*rnf* mutant relative to both the Δ*mmcA* mutant and WWM60, whereas genes in red are uniquely upregulated in the Δ*mmcA* mutant compared to the Δ*rnf* mutant. The seven remaining genes of the Rnf complex in the Δ*mmcA* mutant are shaded in maroon. **b)** Chromosomal organization of the thirteen-gene *fpo* operon in *M. acetivorans*, and the additional *fpoO2* and *fpoF* genes. Double vertical lines indicate genes are located more than 3 kbp away in the genome. The log_2_-fold change in transcript abundance for each gene in the Fpo complex for all pairwise comparisons between the Δ*mmcA* mutant, the Δ*rnf* mutant, and WWM60 are shown in the bar graph. The dashed line on the plot delineates the cutoff for ‘significant’ log_2_-fold change in transcript abundance. In the bar graph: light blue bars represent the expression in the Δ*mmcA* mutant compared to the Δ*rnf* mutant, with higher expression in the Δ*mmcA* mutant denoted by a positive log_2_-fold change. Medium blue bars represent the expression in the Δ*rnf* mutant compared to the parent strain, with higher expression in the Δ*rnf* mutant denoted by a positive log_2_-fold change. Dark blue bars represent the expression in the Δ*mmcA* mutant compared to the parent strain, with higher expression in the Δ*mmcA* strain denoted by a positive log_2_-fold change. **c)** Chromosomal organization of four different methylamine methyltransferase loci. The log_2_-fold change in transcript abundance for each gene at the various loci for all pairwise comparisons between the Δ*mmcA* mutant, the Δ*rnf* mutant, and WWM60 are shown in the bar graphs. The dashed lines on the plots delineate the cutoff for ‘significant’ log_2_-fold change in transcript abundance. In both bar graphs: light green bars represent the expression in the Δ*mmcA* mutant compared to the Δ*rnf* mutant, with higher expression in the Δ*mmcA* mutant denoted by a positive log_2_-fold change. Medium green bars represent the expression in the Δ*rnf* mutant compared to the parent strain, with higher expression in the Δ*rnf* mutant denoted by a positive log_2_-fold change. Dark green bars represent the expression in the Δ*mmcA* mutant compared to the parent strain, with higher expression in the Δ*mmcA* strain denoted by a positive log_2_-fold change.

In addition to the Fpo genes, we also observed significantly higher expression of multiple genetic loci that encode TMA, DMA (dimethylamine), and MMA (monomethylamine) methylamine methyltransferases as well as permeases putatively involved in the transport of these methylated amines in the Δ*rnf* mutant relative to the Δ*mmcA* mutant (Figure 5a, 5c; Supplementary Table 4) (33, 34). Differential expression of these genes could lead to a commensurate change in the rate of transport and conversion of methylated amines to methyl-coenzyme M, an intermediate that feeds into the core methanogenic pathway, which ultimately would affect cell growth as observed in Figure 3. To determine if transcription of these loci was induced in the Δ*rnf* mutant or restricted in the Δ*mmcA* mutant, we compared the expression of these genes in each mutant relative to WWM60 independently. While we did not detect any significant change in transcript levels of any of these genes in the Δ*rnf* mutant relative to WWM60, whereas most of these genes had 4 to 5-fold higher expression in WWM60 relative to the Δ*mmcA* mutant (Supplementary Table 4). These data strongly suggest that the methyltransferase loci are downregulated only when the *mmcA* locus is deleted but not when the entire the entire *rnf* locus is deleted. While the mechanistic details of this regulatory process are beyond the scope of this work, these data bring to light previously unknown global mechanisms in methanogens that coordinate the expression of genes involved in energy conservation (such as *mmcA* and *rnf)* with genes involved in carbon metabolism (such as the *mttCB, mtbCB, mtmCB* methyltransferases involved in growth on TMA, DMA, and MMA, respectively) (33, 34).

Nearly forty genes had significantly higher expression in the Δ*mmcA* mutant relative to the Δ*rnf* mutant and these could be divided into three categories: a) genes that were globally upregulated in the Δ*mmcA* mutant, i.e. these genes were also expressed to a higher level in the Δ*mmcA* mutant relative to WWM60, b) genes that were globally downregulated in the Δ*rnf* mutant i.e. genes that also had a lower expression in the Δ*rnf* mutant compared to WWM60, and c) genes that were only expressed to a higher degree in the Δ*mmcA* mutant relative to the Δ*rnf* mutant, i.e. genes that were not differentially expressed when comparing either mutant to WWM60 (Supplementary Table 5). Of the six genes in the first category, four lacked any recognizable motifs or domains and the other two encode proteins involved in the biosynthesis of asparagine (*asnB*) and post-translational modification of proteins (O-linked N-acetylglucosamine transferase) (Supplementary Table 5) (35, 36). At present, a connection between MmcA and these proteins remains elusive.

About twenty genes were consistently downregulated in the Δ*rnf* mutant and this list includes *rnfCDGEABX* (which validates our methods for identifying changes in transcription), biosynthetic genes, and several genes likely involved in transcriptional regulation (Supplementary Table 5). Downregulated biosynthetic genes included *thiC* (MA4010), a UbiA prenyltransferase domain containing protein, which catalyzes the synthesis of lipophilic compounds that serve as electron carriers in the ETC (37, 38). We hypothesize *ubiA* is involved in the biosynthesis of MP and, in the absence of Rnf, transcription is reduced to modulate levels of MP in the membrane. This hypothesis agrees with a previous study where *ubiA* and other predicted ubiquinone/menaquinone biosynthetic genes were proposed to be involved in MP synthesis and more highly expressed in *M. barkeri* during direct interspecies electron transfer (DIET) to facilitate extracellular electron transport through the membrane (38). Other downregulated biosynthetic genes included a putative operon with an acyl carrier protein and a long chain fatty acyl CoA ligase (MA1027-MA1029). We also observed downregulation of several regulatory genes included an ArsR family transcriptional regulator (MA0504), a response regulator (MA4671), as well as a protein with a DNA-binding helix-turn-helix motif (MA4484) (Supplementary Table 5). The targets of the various regulatory proteins are unknown but might be one of biosynthetic genes mentioned above or the DMS specific methyltransferases, based on the shortened lag time observed in Figure 3d.

Finally, a set of thirteen genes had higher expression only in the Δ*mmcA* mutant relative to the Δ*rnf* strain but were not differentially expressed in comparison to WWM60 (Supplementary Table 5). This list includes signaling proteins like a sensory transduction histidine kinase (MA2256), regulatory genes such as *nikR*, a nickel-responsive transcriptional regulator that controls the expression of nickel-containing enzymes, and cofactor biosynthetic genes like *nadE,* which encodes NAD synthetase that catalyzes the last step in NAD biosynthesis (39, 40). It is tempting to speculate that the signaling proteins or the regulators identified above are linked to the downregulation of the methylamine specific methyltransferases in the Δ*mmcA* strain (Figure 5c; Supplementary Table 4), however detailed mechanistic analyses would be needed to bolster this observation in future work.

Overall, our RNA-sequencing analyses provided clear insights into the genetic basis of the phenotypic distinctions between the Δ*mmcA* strain and the Δ*rnf* strain observed in Figure 3. Downregulation of substrate specific methyltransferases and lower levels of the “headless” Fpo complex, which can potentially regenerate reduced ferredoxin, leads to a more severe growth defect for the Δ*mmcA* mutant relative to the Δ*rnf* mutant on methylated substrates.

### Novel routes for generating a Na^+^ ion gradient and regenerating ferredoxin enable methylotrophic growth in the absence of a functional Rnf complex

To understand how the Δ*rnf* and Δ*mmcA* mutants sustain growth on methylated compounds and the physiological basis for the fitness defect they incur (Figure 3), we compared the transcriptomic profile of each mutant in mid-exponential phase on TMA to that of WWM60 under the same growth conditions (Figure 6a and Figure 6b). Genes were classified as being “differentially expressed” if they met the same criteria listed above. The expression level of genes in WWM60 was considered to be the baseline, so genes with significantly higher or lower transcript levels in the mutants were considered to be upregulated or downregulated respectively.

Only thirteen genes were upregulated and twenty-four genes were downregulated in both mutants compared to WWM60 (Supplementary Table 6). The upregulated genes belonged to three distinct gene clusters (Figure 6a and Figure 6b). First, with the exception of *pstS,* all other genes of the high-affinity phosphate (Pi) transport system (*pstSCAB-phoU)* and alkaline phosphatase (*phoA*) were upregulated between 4.0 to 12-fold in the mutants (Supplementary Table 6) (41). Previous studies with *M. mazei* have shown that cells experiencing Pi starvation upregulate *pstSCAB-phoU* as well as *phoA* (41). Here, we anticipate that these mutants have upregulated the phosphate transport and hydrolysis genes to meet an increased cellular demand for Pi despite slower growth. We suspect that the excess Pi might be needed for ATP synthesis or biosynthesis of methanogenic cofactors, such as tetrahydrosarcinopterin (H_4_SPT) and coenzyme B. The *M. acetivorans* genome also encodes a low-affinity Pi transport system (MA2934-MA2935) that was not differentially expressed in these strains (Supplementary Table 7) (42). Next, several genes in the operon encoding the F_1_F_O_ ATP synthase were upregulated by 4.0 to 8.0-fold in both mutants (Figure 6c; Supplementary Table 6). *M. acetivorans* encodes two different ATP synthases: an archaeal A_1_A_O_ ATP synthase (MA4152-MA4160), which can translocate H^+^ ions and Na^+^ ions concomitantly, that is essential for growth (43), as well as a bacterial F_1_F_O_ ATP synthase (MA2433-MA2441) that is dispensable during methylotrophic methanogenesis and is hypothesized to only translocate Na^+^ ions (44). During methylotrophic growth, the Rnf complex generates a Na^+^ gradient that is used for: a) ATP synthesis by the promiscuous A_1_A_O_ ATP synthase and b) the endergonic transfer of the methyl group from methyl-CoM to H_4_SPT catalyzed by N*^5^*-methyl-H_4_SPT (CH_3_-H_4_SPT): Coenzyme M (CoM) methyltransferase (Mtr) (Supplemental Figure 1) (3, 43). In the absence of a functional Rnf complex, we hypothesize that the A_1_A_O_ ATP synthase relies solely on the proton gradient generated by Fpo and HdrDE for ATP synthesis and the F_1_F_O_ ATP synthase is upregulated to generate a Na^+^ gradient via ATP hydrolysis, which can then be used for the endergonic reaction catalyzed by Mtr. Our hypothesis is further corroborated by the observation that the upregulated genes in the F_1_F_O_ ATP synthase locus are either involved in the assembly of the membrane-embedded F_O_ domain (*atpI*; MA2439) or encode the ‘a’ subunit (*atpB*; MA2437) and the ‘c’ ring (*atpE*; MA2436) of the F_O_ complex that bind and translocate Na^+^ ions (44, 45). In a previous study, a multi-subunit Na^+^/H^+^ antiporter (encoded by the Mrp locus) was shown to play an important role in coupling growth and methanogenesis by generating an optimal Na^+^/H^+^ gradient for efficient ATP synthesis on acetate (46). We did not observe a significant change in the expression of the Mrp locus in either mutant (Supplementary Table 7). Thus, the putative role of the F_O_F_1_ synthase in generating a Na^+^ gradient in the Δ*rnf* and the Δ*mmcA* mutants is non-overlapping with the cellular function of Mrp and other Na^+^/H^+^ antiporters, and potentially highlights different strategies for balancing ion gradients depending on the methanogenic substrate. Finally, the *mtpCAP* locus was upregulated by 8 to 11-fold in the Δ*rnf* mutant and by 16 to 22-fold in the Δ*mmcA* mutant (Supplementary Table 6). Recent studies have shown that the *mtpCAP* locus is involved in the transport and catabolism of the methylated sulfur compound methylmercaptopropionate (MMPA) in *M. acetivorans* (47), however no MMPA was present in our growth media. At present, neither the cause for upregulation of the *mtpCAP* methyltransferase system nor its effect on the physiology of the mutants is clear. However, this observation demonstrates the intricate coupling between methyltransferase regulation and energy conservation in methanogens. Genes that are downregulated in the two mutants have core housekeeping functions and we anticipate that the differential expression of these loci is a consequence of slower growth in the mutants (Supplementary Table 6).

**Figure 6.**
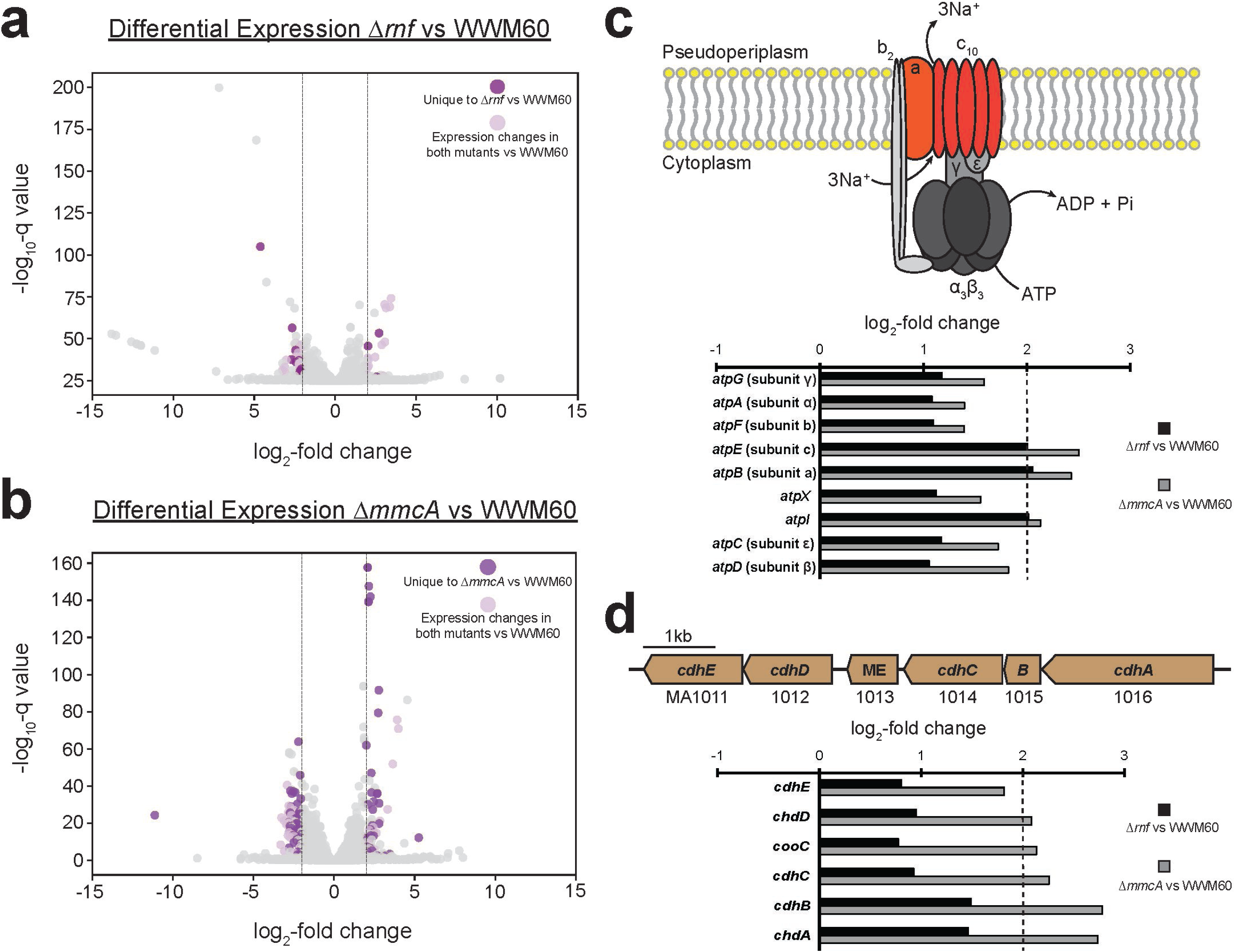
**a)** Volcano plot showing the differential expression of genes between the Δ*rnf* and the parent strain (WWM60). Genes with higher expression in the Δ*rnf* mutant have a positive log_2_-fold change value, while genes with higher expression in the parent have a negative log_2_-fold change value. Dashed lines on the plot delineate the cutoff for ‘significant’ log_2_-fold change in transcript abundance in either strain. Genes in light purple are significantly differentially expressed in both the Δ*rnf* and Δ*mmcA* mutant relative to the WWM60. Genes in dark magenta have significant differential expression only in the Δ*rnf* mutant relative to WWM60. **b)** Volcano plot showing the differential expression of genes between the Δ*mmcA* and the parent strain (WWM60). Genes with higher expression in the Δ*mmcA* mutant have a positive log_2_-fold change value, while genes with higher expression in the parent have a negative log_2_-fold change value. Dashed lines on the plot delineate the cutoff for ‘significant’ log_2_-fold change in transcript abundance in either strain. Genes in light purple are significantly differentially expressed in both the Δ*rnf* and Δ*mmcA* mutant relative to the parent strain. Genes in dark purple have significant differential expression only in the Δ*mmcA* mutant relative to the parent. **c)** Schematic of the Na^+^- translocating F1F0 ATP synthase in *M. acetivorans*. ATP hydrolysis results in the translocation of three sodium ions across the cell membrane to maintain the sodium gradient in the absence of a fully functional Rnf complex (44). Subunits with significant fold change are shaded in orange. The log_2_-fold change in transcript abundance for every gene in the F1F0 ATP synthase between the Δ*rnf* mutant (black bars) and the Δ*mmcA* mutant (gray bars) compared to WWM60 are shown in the bar graph. The dashed line on the plot delineates the cutoff for a ‘significant’ log_2_- fold change in transcript abundance. **d)** Chromosomal organization of the *cdh1* operon in *M. acetivorans*. The log_2_-fold change in transcript abundance for each gene in the *cdh1* operon between the Δ*rnf* mutant (black bars) and the Δ*mmcA* mutant (gray bars) compared to WWM60 are shown in the bar graph. The dashed line on the plot delineates the cutoff for a ‘significant’ log_2_-fold change in transcript abundance.

Next, we analyzed the subset of genes that were differentially expressed in only one of the two mutants. In the Δ*rnf* mutant, three genes were upregulated and ten genes were downregulated in comparison to WWM60 (Supplementary Table 8). Oddly, the *hypF* gene involved in the maturation of hydrogenases (48) was upregulated 6.5-fold, even though *M. acetivorans* lacks any detectable hydrogenase activity during methylotrophic growth conditions (13). In addition, the *kefC* locus encoding a putative glutathione regulated K^+^ efflux system was also upregulated 4-fold in the Δ*rnf* mutant. KefC has been shown to transport Li^+^ and Na^+^ ions (49), therefore we hypothesize that this locus might also aid in establishing a Na^+^ ion gradient in the absence of Rnf. Apart from some genes encoding the pseudo periplasmic substrate binding protein of various ABC transporters, the majority of genes uniquely downregulated in the Δ*rnf* mutant compared to WWM60 did not have a recognizable functional motif (Supplementary Table 8).

In contrast to the Δ*rnf* mutant, a substantially larger number of genes are uniquely induced or suppressed in the Δ*mmcA* mutant (Supplementary Table 9). Among the genes that are upregulated, a few loci are particularly notable. First, genes encoding the carbon monoxide dehydrogenase/acetyl CoA synthase (CODH/ACS) enzyme are upregulated by 4.0 to 7.0-fold. *M. acetivorans* contains two isoforms of CODH/ACS that are encoded by the *cdh1* and *cdh2* operons (Figure 6d) (50). Our initial analysis indicated that five subunits of *cdh1* (*cdhA1*, *cdhB1*, *cdhC1*, *cooC1*, and *cdhD1*) and three subunits of *cdh2* (*cdhC2*, *cooC2*, *cdhD2*) are upregulated in the Δ*mmcA* mutant. Upon closer inspection, we noticed that the two *cdhA* and *cdhB* homologs are relatively divergent at the sequence level (∼80% amino acid identity) whereas the *cdhC*, *cooC*, *cdhD* homologs share >97% amino acid identity (14). Thus, the upregulation of the latter set in *cdh2* could be an artefact of the RNA-sequencing analysis pipeline, similar to previous reports of transcript cross-reactivity between *cdh* operons in *M. mazei* (51). Regardless, both isoforms are known to be functionally redundant and can catalyze the catabolism of acetate during acetoclastic methanogenesis or anabolic acetyl-CoA synthesis by the Wood-Ljundahl (WL) pathway during methylotrophic growth (50). Since carbon fixation by the WL pathway requires reduced ferredoxin, increased expression of CODH/ACS could serve as an alternate route for regenerating reduced ferredoxin in the absence of a functional Rnf complex in the Δ*mmcA* mutant. A 4.0-fold increase in expression of the regulatory protein encoded by the *mreA* locus in the Δ*mmcA* mutant is likely linked to the upregulation of *cdh1*. MreA is a global regulator of methanogenic pathways in *M. acetivorans* and has been shown to activate genes important for acetoclastic methanogenesis (such as *cdh1*) and to repress transcription of genes that play a crucial role in methylotrophic methanogenesis (including the methylamine-specific methyltransferases and the *fpo* locus) (Figure 5b and Figure 5c) (52). In a previous study, the *ack/pta* locus encoding acetate kinase and phosphate acetyltransferase were downregulated in a Δ*mreA* strain, however we did not observe any significant change in the expression of these loci in the Δ*mmcA* mutant (Supplementary Table 7) (52). Thus, it is likely that other regulators in addition to MreA are also involved in the upregulation of genes required for the catabolism of acetate. We also observed a 5.6-fold increase in expression of a gene encoding a bile acid: Na^+^ symporter family protein (MA2632) in the Δ*mmcA* mutant, which may be linked to the maintenance of the Na^+^ ion gradient across the membrane. Among the genes that were significantly downregulated in the Δ*mmcA* mutant, two loci are of particular interest. First, the *pylBCD* locus involved the biosynthesis of the 22^nd^ amino acid, pyrrolysine (Pyl), is downregulated by 4.0 to 4.6-fold (53). Whether the *pyl* genes are a part of the same regulon as the concomitantly downregulated methylamine methyltransferases (Figure 5c) or if the expression of the *pyl* genes is controlled by the amount of the methylamine methyltransferases remains unclear. Next, two genes (*cfbA* and *cfbE)* involved in the biosynthesis of Factor 430 (F_430_), a Ni-containing cofactor associated with MCR, were downregulated significantly (54, 55). Downregulation of F_430_ production might free up more Ni for increased production of the Ni-containing CODH/ACS enzyme in Δ*mmcA* mutant.

## Conclusions

The acquisition of an ETC likely spurred rampant ecological diversification in members of the Order *Methanosarcinales.* For several decades, the bioenergetic complexes that comprise the ETC in these archaea were studied in isolation using *in vitro* techniques. While these studies have provided substantial insights into the biochemical mechanisms that facilitate electron transfer reactions, an *in vivo* perspective on the ETC and its interplay with metabolism has been lacking. In this study, we performed comprehensive genetic, phenotypic, and transcriptomic analyses of *M. acetivorans* mutants that either lack the entire Rnf bioenergetic complex or a just a single subunit encoding an MHC called MmcA. Our growth analyses are congruent with a previous study, which also demonstrated that Rnf complex is essential for growth on acetate but not on methylated compounds (15). Our transcriptomic analyses provide evidence of potential alternative mechanisms for ferredoxin regeneration in each mutant (Figure 5b and Figure 6b). In the Δ*rnf* mutant, a “headless” Fpo complex might serve as a new entry point for electrons from ferredoxin (Figure 5b; Supplementary Figure 2) in the ETC (32), whereas acetyl CoA synthesis mediated by CODH/ACS could possibly regenerate the reduced ferredoxin pool the Δ*mmcA* mutant (Figure 6b) (26). The “headless” Fpo backup strategy is coupled to proton translocation and would theoretically conserve more energy for the Δ*rnf* mutant compared to the CODH/ACS strategy in the Δ*mmcA* mutant. Accordingly, the Δ*rnf* mutant has faster growth rate than then Δ*mmcA* mutants on methylated compounds (Figure 3). We anticipate that the potential alternative strategies for ferredoxin regeneration stem from distinct regulatory responses by the cell to the loss of either MmcA or the entire Rnf complex. Our transcriptomic data corroborates this hypothesis, in which we saw upregulation of the global methanogenesis protein MreA in the Δ*mmcA* mutant but not the Δ*rnf* mutant (Figure 6; Supplementary Table 9). Higher expression of MreA would lower the expression of *fpo* locus in the Δ*mmcA* mutant during methylotrophic methanogenesis (52). Similarly, induction of MreA during acetoclastic growth might also explain the lethal phenotype for both mutants (52). These data showcase the sheer diversity of energy conservation strategies present in *M. acetivorans*, and likely other members of the *Methanosarcinales*, which enable these organisms to thrive in a wide array of ecological niches.

Additionally, based on our phenotypic (Figure 3 and Figure 4) and transcriptomic analyses (Figures 5 and Figure 6), we observe that the impact of deleting the Rnf complex or MmcA extends far beyond energy conservation in the cell. Genes involved in Na^+^ ion transport, amino acid biosynthetic pathways, substrate specific methyltransferases for methylotrophic methanogenesis, transcriptional regulators, and many other loci were differentially expressed in one or both mutants (Figure 5 and Figure 6). These dramatic transcriptional changes underscore a complex and intricate regulatory network that connects carbon transformation by methyltransferases and energy conservation during methanogenic growth. Further analyses of regulatory genes identified in this work and similar studies with other components of the ETC will ultimately provide systems-level insights into methanogenesis in *Methanosarcina acetivorans,* and will deepen our understanding of these ecologically-relevant microbes.

## Materials and Methods

### Media and culture conditions

All *Methanosarcina* strains were grown at 37°C without shaking in bicarbonate-buffered high-salt (HS) liquid medium containing either 50 mM trimethylamine hydrochloride (TMA), 125 mM methanol, 40 mM sodium acetate, or 20 mM dimethylsulfide (DMS) as a growth substrate (56). TMA, methanol, and acetate were added prior to autoclaving whereas DMS was added after autoclaving from a 200 mM stock solution prepared in HS medium with no other carbon sources. For mutant generation, the growth medium contained 50 mM TMA as the growth substrate and agar solidified HS + TMA media was obtained by adding 1.5% w/v agar (Sigma-Aldrich, St. Louis, MO, USA). To select for transformants, puromycin (Pur) (RPI, Mount Prospect, IL, USA) was added to HS + TMA agar medium before solidification to a final concentration of 2 ug/mL from a 1000X sterile, anaerobic stock solution with N_2_ gas in the headspace at 55-69 kPa. HS + TMA + Pur agar plates were incubated at 37°C in an intra-chamber anaerobic incubator with N_2_/CO_2_/H_2_S (79.9%/20%/0.1%) in the headspace, as described previously (57). All *Escherichia coli* strains were grown in Lysogeny broth (LB) at 37°C in a shaking incubator (Thermo Fisher Scientific, Waltham, MA, USA) at 250 rpm. To select for the desired plasmids, antibiotics were added to cultures to final concentrations of 25 µg/mL for kanamycin, and/or 10 µg/mL for chloramphenicol as listed in Supplementary Table 10. For plasmid extraction, rhamnose was added to a final concentration of 10 mM to *E. coli* cultures prior to incubation to increase the plasmid copy number of pDN201- and pJK029A- derived plasmids.

### Construction of Methanosarcina acetivorans mutants

Liposome-mediated transformation of *M. acetivorans* was performed as previously described (58). Briefly, 20 mL of late-exponential phase (∼0.8 OD_600_) cultures growing on HS + TMA were harvested by centrifugation, the supernatant was decanted, and the cell pellet was resuspended in 1 mL of anaerobic bicarbonate-buffered, isotonic sucrose (pH = 7.4) containing 100 µM cysteine. Next, 25 uL of N-[1-(2,3-Dioleoyloxy)propyl]-N,N,N-trimethylammonium methylsulfate (DOTAP) (Roche Diagnostics Deutschland GmbH, Mannheim, Germany) and 2 ug of plasmid DNA were incubated for 30 mins in 75 uL of buffered, isotonic sucrose to allow for DNA uptake into liposomes. After incubation, the DOTAP + DNA mixture was added in full to the cell suspensions. Suspensions of cells + DOTAP + DNA were incubated for 4 hours at room temperature in an anaerobic chamber with CO_2_/H_2_/N_2_ (10/5/85) in the headspace before inoculation into 10 mL of HS + TMA. Outgrowths of transformed cells were incubated at 37°C for 12-16 hours before plating on agar-solidified HS + Pur + TMA using a sterile spreader (56). Plasmids used for mutant generation are described in Supplementary Table 10. Mutant colonies were genotyped using primers detailed in Supplementary Table 11, and a full list of strains used in this study is provided in Supplementary Table 12.

### Growth Assays for Methanosarcina acetivorans mutants

For growth analysis, 11 mL cultures were grown at 37°C without shaking (HeraTherm™General Protocol Microbiological Incubator, Thermo Fisher Scientific, Waltham, MA, USA) in Balch tubes with N_2_/CO_2_ (80/20) at 55-69 kPa in the headspace. Growth of three independent biological replicates was measured by determining the optical density of cultures at 600nm (OD_600_) using a UV-Vis Spectrophotometer (Gensys 50, Thermo Fisher Scientific, Waltham, MA, USA). A Balch tube containing 10 mL of HS medium with the appropriate growth substrate was used as a ‘blank’ for optical density measurements. For growth on TMA or methanol, cells were acclimated to the growth substrate for a minimum of four generations prior to quantitative growth measurements. Growth measurements on acetate and DMS were performed with cells previously grown on TMA.

Approximately 1 mL of late exponential phase culture was harvested and served as the inoculum into 10 mL of fresh medium for growth analyses. Growth data were log10-transformed and plotted versus time. A linear regression was fitted to the data to include at least 5 data points on the growth curve for a regression coefficient (R^2^) ≥ 0.97. Growth rate (gr) was calculated as the slope of the linear fit multiplied by 2.303, and the doubling time was calculated as T_d_ = 0.6932/gr. Lag time was calculated by subtracting the Y-intercept value from the log10- transformed initial OD_600_ reading and dividing by the slope of the linear fit. For maximum OD_600_ measurements, approximately 1 mL of early stationary phase culture was harvested and diluted into 10 mL of fresh HS medium containing the same substrate used for growth. An OD_600_ measurement of the diluted culture was then multiplied by 11 to approximate the maximum OD_600_ value. Growth curve plots, determination of doubling time, lag time, and statistical analyses were obtained using Microsoft Excel Version 16.55. Plots of OD_600_ versus time were generated using GraphPad/Prism 9.3.1.

### DNA extraction and sequencing

Cells from a 10 ml culture of DDN009 (Δ*mmcA*) incubated in in HS + TMA at 37 °C were harvested at late-exponential phase (OD_600_ ∼0.8) for genomic DNA extraction using the Qiagen blood and tissue kit (Qiagen, Hilden, Germany). The concentration of genomic DNA was measured using a Nanodrop One Microvolume UV-Vis Spectrophotmeter (Thermo Scientific, Waltham, MA, USA). Genomic DNA was shipped to the Microbial Genome Sequencing Center, Pittsburgh, PA, USA, where sequencing libraries preparation and sequencing was performed. Sequencing reads were aligned to the *Methanosarcina acetivorans* C2A genome and mutations were identified using Breseq version 0.35.5. Illumina sequencing reads for DDN009 have been deposited to the Sequencing Reads Archive (SRA) and the BioProject accession number will be made available upon publication.

### RNA extraction and sequencing

WWM60 (parent), WWM1015 (Δ*rnf*) and DDN009 (Δ*mmcA*) pre-acclimated on TMA were inoculated in quadruplicate from late exponential phase cultures (OD_600_ ∼0.8) into 10mL of fresh HS + TMA in Balch tubes with N_2_/CO_2_ (80/20) at 55-69 kPa in the headspace and grown at 37°C without shaking (Isotemp™ Microbiological Incubator, Thermo Fisher Scientific, Waltham, MA, USA). One Balch tube was used to monitor growth as a proxy for the other replicates by measuring the OD_600_ routinely using a UV-Vis Spectrophotometer (Gensys 50, Thermo Fisher Scientific, Waltham, MA, USA). Once the measured OD_600_ reached approximately ½ maximum value (0.750-0.850), RNA was harvested from the remaining three culture tubes. For RNA extraction, 1 mL of culture was added to 1 mL of Trizol pre-warmed to 37°C (Life Technologies, Carlsbad, CA, USA) and incubated at room temperature for 5 minutes. Next, 2 mL of 100% ethanol was added to each sample and RNA extraction was performed using the Qiagen RNeasy Mini Kit (Qiagen, Hilden, Germany) according to the manufacturer’s instructions. The concentration and quality of RNA samples was determined using a Nanodrop One/One^C^ UV Spectrophotometer (Thermo Fisher Scientific, Waltham, MA, USA) before storage at −80°C. Samples were submitted to the Microbial Genome Sequencing Center (Pittsburg, PA) for DNase treatment, rRNA depletion, library preparation, and Illumina paired-end sequencing. On average 98% of reads were mapped to the *M. acetivorans* C2A genome. Raw transcript reads were deposited to the Sequence Read Archive (SRA) and the BioProject accession number will be made available upon publication.

### RNAseq analysis

Reads in FASTQ format were uploaded to KBase in a new ‘Narrative’: a Jupyter-based user interface in which the raw reads are processed using ‘apps’ and the output/result of each app is recorded and accessible for download (59). Raw transcript reads were grouped by strain in a ‘SampleSet’ using the app ‘Create RNA-seq SampleSet’. Next, the SampleSet was selected as input for read alignment using the HISAT2 (v.2.1.0) app with the *Methanosarcina acetivorans* C2A genome serving as the reference for mapping. Aligned reads were assembled using the Cufflinks (v2.2.1) app with the *M. acetivorans* C2A genome input as the reference. The output of Cufflinks was exported an Expression Set and individual expression matrices for each strain and associated replicates. The expression values for each gene were reported as log_2_(FPKM) (fragments per kilobase per million mapped reads). Finally, differential expression matrices for each pairwise combination of strains were generated by uploading the log_2_(FPKM) Expression Set to the DESeq2 (v1.20.0) app. Changes in transcript abundance were considered significant between strains if a ≥±2 log_2_-fold (q-values ≤ 0.05) change was seen. Differential expression of genes was visualized in volcano plots constructed using a Python script and a table with the locus tag, log_2_-fold change in expression, q-value, and assigned color for each gene. Before plotting, select genes were highlighted in colors other than gray by manual curation of the table in Microsoft Excel (version 16.62), and genes with a q-value of zero were assigned a value of 1E-300. Volcano plots show log_2_-fold change on the x-axis and log_10_(q-value) on the y-axis for each gene.

## Acknowledgements

The authors would like to acknowledge Katie Shalvarjian for assistance with data visualization and other members of the Nayak lab for their feedback and input.

